# The ‘C3aR antagonist’ SB290157 is a partial C5aR2 agonist

**DOI:** 10.1101/2020.08.01.232090

**Authors:** Xaria X. Li, Vinod Kumar, John D. Lee, Trent M. Woodruff

## Abstract

Innate immune complement activation generates the C3 and C5 protein cleavage products C3a and C5a, defined classically as anaphylatoxins. C3a activates C3a receptors (C3aR), while C5a activates two receptors (C5aR1 and C5aR2) to exert their immunomodulatory activities. The non-peptide compound, SB290157, was originally reported in 2001 as the first C3aR antagonist. In 2005, the first report on non-selective nature of SB290157 was published, where the compound exerted clear agonistic, not antagonistic, activity in variety of cells. Other studies also documented non-selective activities of this drug *in vivo*. These findings severely hamper data interpretation regarding C3aR when using this compound. Unfortunately, given the dearth of C3aR inhibitors, SB290157 still remains widely used to explore C3aR biology (>70 publications to date). Given these issues, in the present study we aimed to further explore SB290157’s pharmacological selectivity by screening the drug against three human anaphylatoxin receptors, C3aR, C5aR1 and C5aR2, using transfected cells. We first confirmed that SB290157 acts as a potent agonist at human C3aR. We also identified that SB290157 exerts partial agonist activity at C5aR2 by mediating β-arrestin recruitment at higher compound doses. Notably, SB290157’s activity at C5aR2 was more potent and efficacious than the current ‘lead’ C5aR2 agonist P32. Notwithstanding this, SB290157 showed inhibitory effect on C3a-mediated signalling in primary human macrophages. Our results therefore provide even more caution against using SB290157 as a research tool to explore C3aR function. Given the reported immunomodulatory and anti-inflammatory activities of C3aR and C5aR2 agonism, any function observed with SB290157 could be due to these off target activities.

## 1 INTRODUCTION

The complement system is an essential component of innate immunity, responding to external and internal insults. Complement activation, through all pathways including the classical, lectin and alternative pathways, generates the cleavage complement peptides C3a and C5a, which serve as important innate effectors, modulating multiple aspects of immune cell function (1). C3a is one of the most abundant complement activation products generated from the cleavage of C3 by C3 convertases (2). Human C3a, consisting of 77 amino acids (3), interacts potently with the canonical 7-transmembrane G-protein coupled receptor, C3a receptor (C3aR), which is widely expressed by all myeloid cells (such as neutrophils and monocytes), activated T lymphocytes and B lymphocytes (4–8), and also non-leukocytes such as microglia, astrocytes and endothelial cells (4). C3aR is predominantly coupled to the pertussis toxin-sensitive G protein Gα_i2_ (9). Upon activation, C3aR dampens the intracellular cAMP/PKA signalling, induces the PI3K/Akt pathway, intracellular calcium mobilisation and ERK1/2 phosphorylation, and recruits β-arrestins (4). The nature of C3a action appears to be rather nuanced, with both pro- and anti-inflammatory responses reported in the literature (2).

The non-peptide small molecule SB290157 was first reported in 2001 (10). Functioning as a competitive C3aR antagonist (10), SB290157 has served as a major pharmacological tool used to explore C3aR biology. For instance, SB290157 treatment was shown to improve survival in experimental lupus nephritis (11), ameliorate anti-ovalbumin polyclonal antibody–induced arthritis (12), reduce infarct volume in a model of thromboembolic stroke (13), and rescue cognitive impairment in a model of Alzheimer’s disease (14). The promising results shown by administering SB290157 in these experimental models has heightened interest in C3aR blockade as a therapeutic strategy for diverse diseases. However, despite the promising initial preclinical efficacy of SB290157, studies performed in the mid 2000’s demonstrated that this compound possesses clear agonistic (not antagonistic) activities on C3aR, in cells with high levels of C3aR expression (15). Furthermore, independent studies document undefined off-target effects when SB290157 is administered *in vivo*, even at relatively low doses (16, 17). Likely due to these non-selective activities, SB290157 has not progressed commercially as a therapeutic. However, despite these on- and off-target limitations, SB290157 continues to be used widely in the field to block C3aR activity. Indeed, to the best of our knowledge, over 70 research articles have used this drug to explore roles for C3aR, presumably in part due to the limited availability of alternate therapeutic approaches.

Human C3aR shares 38 % (116/309 residues) similarity with human C5aR1 (4, 18), and a 55% similarity in the transmembrane domains with C5aR2 (19). Due to the close homology between C3a and C5a receptors, ligands designed to target one of these receptors may have promiscuous actions on the other two (20, 21). SB290157 was previously shown to be devoid of C5aR1-antagonistic properties in human neutrophils or C5aR1-expressing rat basophilic leukemia cells (10). However, to date, whether SB290157 exerts activity on C5aR2 has not been reported. In the present study, we thus aimed to further characterise the pharmacological activity of SB290157 on all three anaphylatoxin receptors, C3aR, C5aR1 and C5aR2, by employing various signalling assays in transfected cell lines and primary human macrophages. We identified that in addition to acting as a C3aR agonist on transfected cells, SB290157 is also a partial agonist for C5aR2, further highlighting caution against utilising this drug in experimental studies examining C3aR biology.

## 2 MATERIALS AND METHODS

### 2.1 Ligands and materials

The C3aR inhibitor, SB290157 trifluoroacetate salt, and purified human C3a was purchased from Merck (Perth, Australia). Recombinant human C5a (rhC5a) was purchased from Sino Biological (Beijing, China). Bovine serum albumin (BSA) was purchased from Merck (Perth, Australia). For cell culture, trypsin-EDTA, HBSS, HEPES, Dulbecco’s Modified Eagle’s Medium (DMEM), phenol-red free DMEM, Ham’s F12, Iscove’s Modified Dulbecco’s Medium and Penicillin-Streptomycin were purchased from Thermo Fisher Scientific (Melbourne, Australia). Dulbecco’s phosphate-buffered saline was purchased from Lonza (Melbourne, Australia).

### 2.2 Cell culture

The following cell lines were cultured as previously described (22). Briefly, Chinese hamster ovary cells stably expressing the human C5aR1 (CHO-C5aR1) or human C3aR (CHO-C3aR) were maintained in Ham’s F12 medium containing 10% foetal bovine serum (FBS), 100 IU/ml penicillin, 100 μg/ml streptomycin and 400 μg/ml G418 (InvivoGen, San Diego, USA). Human embryonic kidney 293 cells (HEK293) were maintained in DMEM medium containing 10% FBS, 100 IU/ml penicillin and 100 μg/ml streptomycin. All cell lines were maintained in T175 flasks (37°C, 5% CO_2_) and subcultured at 80-90% confluency using 0.05% trypsin-EDTA in DPBS.

Human monocyte-derived macrophages (HMDM) were generated and cultured as previously described (23, 24). Briefly, human buffy coat blood from anonymous healthy donors was obtained through the Australian Red Cross Blood Service (Brisbane, Australia). Human CD14+ monocytes were isolated from blood using Lymphoprep density centrifugation (STEMCELL, Melbourne, Australia) followed by CD14+ MACS magnetic bead separation (Miltenyi Biotec, Sydney, Australia). The isolated monocytes were differentiated for 7 days in Iscove’s Modified Dulbecco’s Medium supplemented with 10% FBS, 100 IU/ml penicillin, 100 μg/ml streptomycin and 15 ng/ml recombinant human macrophage colony stimulating factor (Lonza, Melbourne, Australia) on 10 mm square dishes (Bio-strategy, Brisbane, Australia). Non-adherent cells were removed by washing with DPBS, and the adherent differentiated HMDMs were harvested by gentle scraping.

### 2.3 Phospho-ERK1/2 assays

Ligand-induced ERK1/2 phosphorylation was assessed using the AlphaLISA *Surefire Ultra* p-ERK1/2 (Thr202/Tyr204) kit (PerkinElmer, Melbourne, Australia) following the manufacturer’s protocol as previously described (23, 24). Briefly, CHO-C3aR, CHO-C5aR1 or HMDMs were seeded (50,000/well) in tissue culture-treated 96-well plates (Corning, USA) for 24 h and serum-starved overnight. All ligand dilutions were prepared in serum-free medium (SFM) containing 0.1% BSA. For inhibition assays, cells were pre-treated with SB290157 for 30 min before agonist addition. For stimulation, cells were treated with ligands at the indicated concentrations for 10 min at RT and then immediately lysed using AlphaLISA lysis buffer on a microplate shaker (450 rpm, 10 min). For the detection of phospho-ERK1/2 content, cell lysate (5 μL/well) was transferred to a 384-well ProxiPlate (PerkinElmer, Melbourne, Australia) and added to the donor and acceptor reaction mix (2.5 μL/ well, respectively), followed by a 2-h incubation at RT in the dark. On a Tecan Spark 20M (Tecan, Männedorf, Switzerland), the plate was measured using standard AlphaLISA settings.

### 2.4 BRET assays measuring β-arrestin 2 recruitment to C5aR2

The C5a-mediated β-arrestin 2 recruitment to C5aR2 was measured using bioluminescence resonance energy transfer (BRET)-based assay as previously described (24, 25). Briefly, HEK293 cells were transiently transfected with β-arrestin 2-*Renilla* luciferase 8 (Rluc8) and human C5aR2-Venus constructs using XTG9 (Roche, Sydney, Australia) for 24 h. Transfected cells were then seeded (100,000/well) onto white 96-well plates (Corning, USA) in phenol-red free DMEM containing 5% FBS overnight. For BRET assay, cells were firstly incubated with the substrate EnduRen (30 μM, Promega, Sydney, Australia) for 2 h (37°C, 5% CO_2_). All ligands were prepared in SFM containing 0.1% BSA. On a Tecan Spark 20M microplate reader (Tecan, Männedorf, Switzerland) (37°C), the BRET light emissions (460-485 and 520-545 nm) were continuously monitored for 25 reads with respective ligands added after the first 5 reads. The ligand-induced BRET ratio was calculated by subtracting the emission ratio of Venus (520-545 nm)/ Rluc8 (460-485 nm) of the vehicle-treated wells from that of the ligand-treated wells.

### 2.5 Data collection, processing and analysis

All experiments were conducted in triplicates and repeated on 3 separate days (for cell lines) or using cells from 3 or more donors (for HMDMs) unless otherwise specified. Data was analysed using GraphPad software (Prism 8.4). Data from each individual repeat was normalised accordingly before being combined and expressed as mean ± standard error of the mean (S.E.M.) unless otherwise described. For all dose-response studies, logarithmic concentration-response curves were plotted using combined data and analysed to determine the respective potency values. Statistical analysis comparing the maximum of C3aR activation between SB290157 and human C3a was conducted using an unpaired Student’s t-test. Differences were deemed significant when *P* < 0.05.

## 3 RESULTS

### 3.1 SB290157 potently activates ERK signalling in CHO-C3aR cells

First, we aimed to confirm the prior reported C3aR-agonistic property of SB290157 on an overexpression cell line (15). We measured C3aR-mediated ERK signalling as a readout in CHO cells stably expressing human C3aR. As expected, SB290157 potently induced phosphorylation of ERK in CHO-C3aR cells in a dose-dependent fashion, with a highly potent EC_50_ of 0.46 nM (**Figure 1A**). Notably, the sub-nanomolar potency of SB290157 is comparable to the 0.39 nM potency demonstrated by human C3a, however, the maximum level of C3aR activation caused by SB290157 was 16% lower than human C3a (P=0.0082), similar to a prior report (15). When SB290157 was counter-screened in CHO cells overexpressing human C5aR1, the ligand did not cause significant activation or inhibition of human C5aR1-mediated phopho-ERK1/2 activity (**Figure 1B, C**). Therefore, we confirmed that SB290157 behaves as potent agonist in C3aR-transfected cells, but is inactive at C5aR1.

**Figure 1.**
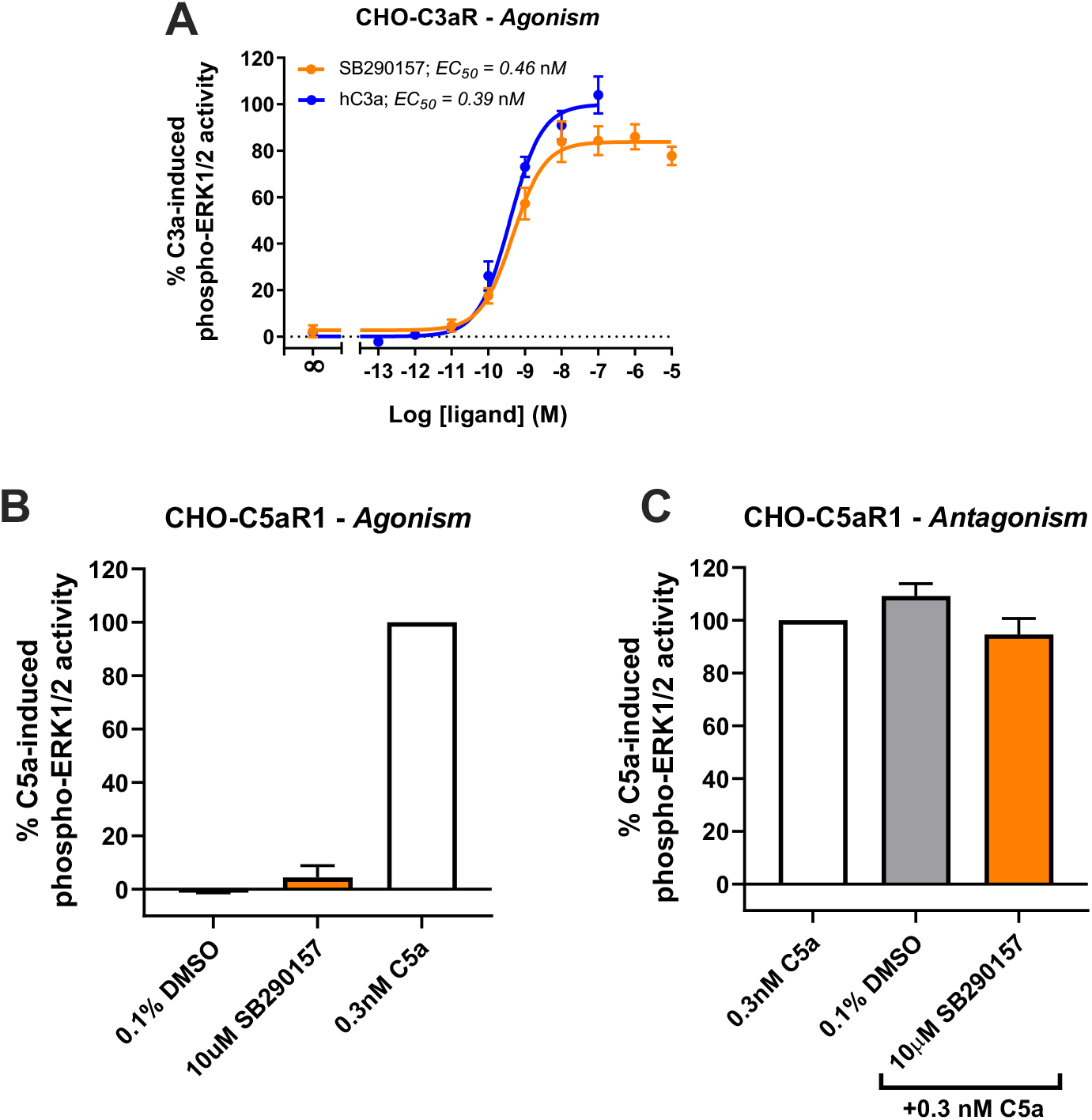
SB290157 potently activates C3aR-mediated ERK signalling in transfected CHO cells. **(A)** Agonism testing of SB290157 on CHO-C3aR cells. Serum-starved CHO-C3aR cells (50,000/well) were stimulated with respective concentrations of SB290157 or purified human C3a for 10 min before being lysed. **(B)** Agonism testing of SB290157 on CHO-C5aR1 cells. Serum-starved CHO-C5aR1 cells (50,000/well) were stimulated with the respective ligands at the indicated concentrations for 10 min before being lysed. **(C)** Antagonism testing of SB290157 on CHO-C5aR1 cells. Serum-starved CHO-C5aR1 cells (50,000/well) were pre-treated with SB290157 (10 μM) or vehicle (0.1% DMSO) for 30 min before being stimulated with 0.3 nM of C5a for 10 min and then lysed. The phospho-ERK1/2 content in the cell lysate was measured and normalised to the maximum C3a-induced (for A) or 0.3 nM C5a-induced (for B, C) levels before being combined. Data represent mean ± S.E.M. of triplicate measurements from 3-6 independent experiments (n = 3-6).

### 3.2 SB290157 possesses off-target activity on human C5aR2

We next examined the potential activity of SB290157 on the other closely related complement receptor, C5aR2. As a non-canonical G-protein coupled receptor, C5aR2 does not couple to the common classes of G proteins and is devoid of the classical G protein-mediated signalling activities (26, 27). Human C5aR2 activation however recruits β-arrestins, which can be used as a readout of receptor activation (25, 28). We thus assessed the potential activity of SB290157 on human C5aR2-β-arrestin 2 interaction using a BRET assay established in HEK293 cells, and compared to that of the existing and widely used C5aR2-selective agonist, P32 (Ac-RHYPYWR-OH) (29).

Surprisingly, SB290157 dose-dependently induced C5aR2-mediated β-arrestin 2 recruitment, with a micromolar potency (EC_50_) of 16.1 μM (**Figure 2A, B**), which is slightly less potent compared to P32 (EC_50_ = 5.9 μM). The maximum level of β-arrestin 2 recruitment to C5aR2 induced by SB290157 however was higher relative to that of P32. We next compared the C5aR2 activity of SB290157 to that of human C5a (**Figure 2C**), and observed that, similar to P32, the C5aR2-agonistic activity of SB290157 is also partial, reaching 31% of the level induced by C5a. Thus, our results clearly show that at higher concentrations, SB290157 also partially activates C5aR2.

**Figure 2.**
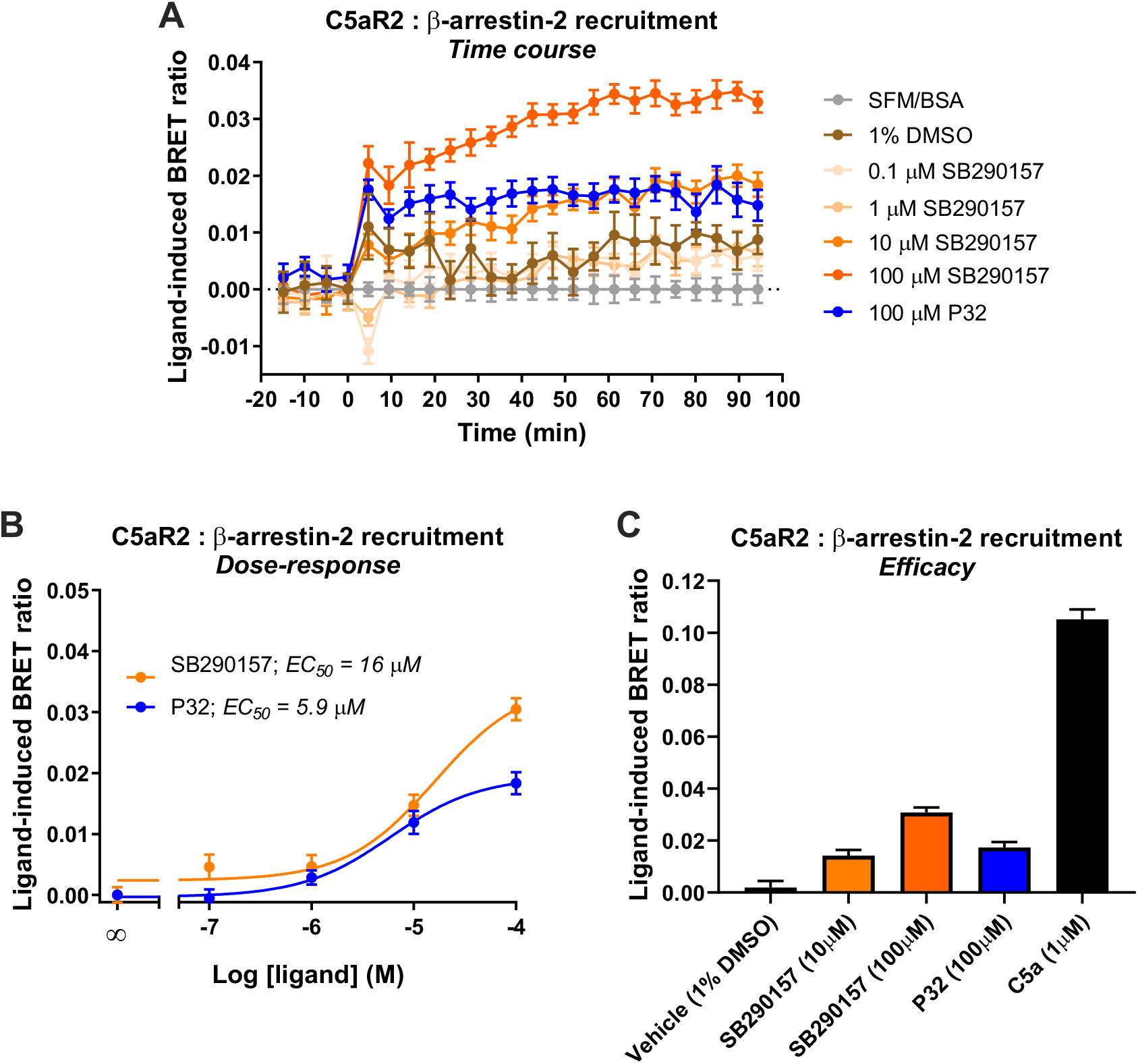
SB290157 induces C5aR2-mediated β-arrestin 2 recruitment in transfected HEK293 cells. HEK293 cells were transiently transfected using C5aR2-Venus and β-arrestin 2-Rluc8 BRET pairs for 24 hours and seeded (100,000/well) overnight. Filtered light emissions between 460-490 nm (Rluc8) and 520-550 nm (Venus) were continually monitored for 90 min with SB290157, P32, C5a or vehicle added at the 0 min time point. Data represent **(A)** the time course of ligand-induced BRET ratios (Venus/Rluc8 emission ratio) caused by the respective ligands at the indicated concentrations, **(B)** the corresponding dose-response curves for SB290157 and P32 at 40 min and post ligand addition, and **(C)** the ligand-induced BRET ratios (efficacy) of SB290157, P32 or C5a at 40 min post ligand addition. Data represent the mean ± S.E.M. of triplicate measurements from 3-5 independent experiments (n = 3-5).

### 3.3 SB290157 potently inhibits C3a-mediated ERK signalling in human monocyte-derived macrophages

We next aimed to determine the activity of SB290157 in primary human immune cells, specifically HMDMs, which is an established and widely used model of resting tissue macrophages that express high levels of human C3aR and the C5a receptors (30, 31). Human C3a potently activate ERK1/2 phosphorylation in HMDMs (23). We therefore assessed the ability of SB290157 to inhibit this C3a-mediated signalling pathway, and found that this compound dose-dependently inhibited C3a-induced ERK signalling in HMDMs (**Figure 3A**). The IC_50_ of SB290157 (236 nM), was ~16-fold more potent than previously reported data (IC_50_ = 3.8 μM), determined using an intracellular calcium mobilisation assay in the same cells (32). The discrepancy could be explained by the competitive nature of SB290157-mediated inhibition. The lower C3a used in the present study, 5 nM versus 100 nM in Rowley et al. (32), would give rise to a lower apparent IC_50_ value of SB290157 (33). In marked contrast to the CHO-C3aR data, SB290157 did not display any agonistic activity on ERK signalling in HMDMs at the highest possible dose tested of 100 μM, although human C3a displayed an EC_50_ of 0.21 nM (**Figure 3B**), in accordance with the previous data (23, 32). We also performed dose titrations of SB290157 to ensure the absence of agonism was not a result of the ERK signalling being down-regulated, as experienced by higher concentrations of human C3a (**Figure 3B**). Thus, at least for in primary human macrophages, SB290157 retains its potent C3a-inhibitory activity.

**Figure 3.**
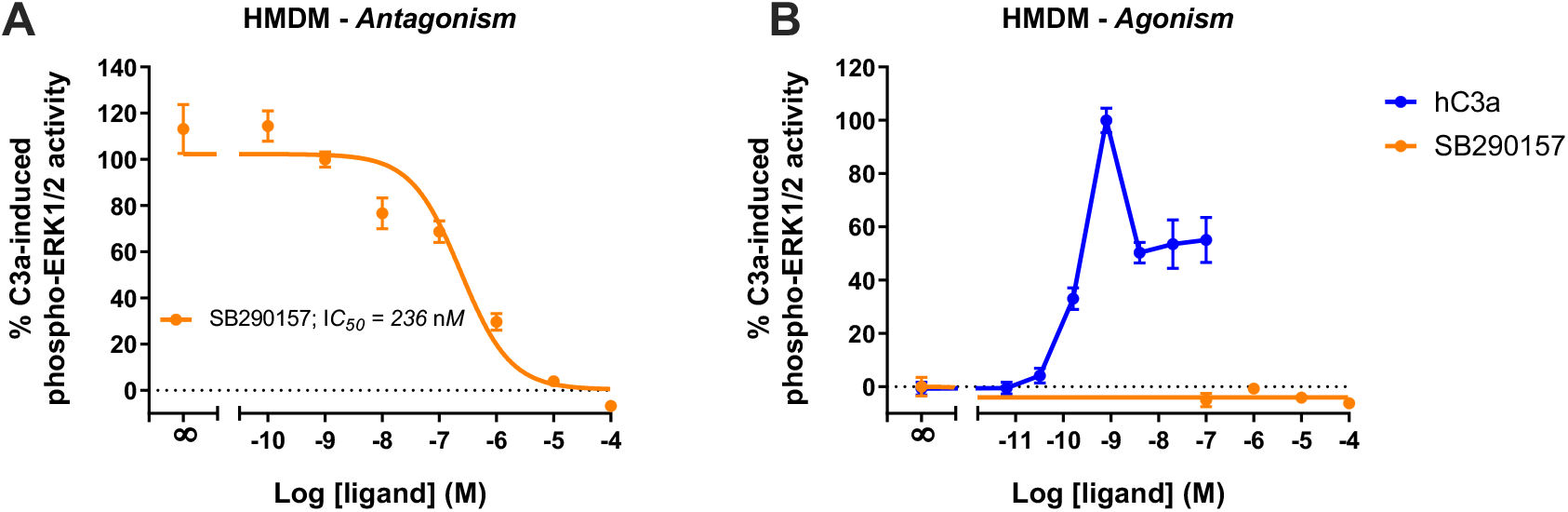
SB290157 potently inhibits C3aR-mediated ERK signalling in human monocyte-derived macrophages. **(A)** Antagonism testing of SB290157 on HMDMs. Serum-starved HMDMs (50,000/well) were pre-treated with various doses of SB290157 for 30 min before being stimulated with purified human C3a (5 nM) for 10 min and then lysed. **(B)** Agonism testing of SB290157 on HMDMs. HMDMs were stimulated with respective doses SB290157 for 10 min and then lysed. The phospho-ERK1/2 content in the cell lysate was measured and normalised to the C3a-induced levels before being combined. Data represent mean ± S.E.M. of triplicate measurements using cells from 2-3 independent donors (n = 2-3).

## 4 DISCUSSION

In this study, we examined the pharmacological properties of the widely-used C3aR antagonist, SB290157 on human C3aR, C5aR1 and C5aR2 receptors, utilising both transfected overexpression cell lines and primary human macrophages. First, in an overexpression system of CHO-C3aR cells, we confirmed the agonistic property of SB290157 on ERK signalling, mediated through C3aR. The sub-nanomolar potency of SB290157 is comparable to that of human C3a, despite SB290157’s much lower affinity on human C3aR compared to C3a (K_i_ = 210 nM for SB290157, versus 0.09 nM for human C3a) (15). To account for this, the high K_i_ of SB290157 mainly results from its relatively large K_off_ rate (15). This property is not reflected well by measuring ERK signalling, a pathway characterised by extensive amplification, such that a transient activation of a small proportion of receptors is sufficient to trigger a complete response (34). Thus, our results indicate that SB290157 may act as a partial agonist with low intrinsic efficacy, which can elicit a near-complete response in cells with high receptor density (15, 35).

Next, when counter-screening against a closely relative complement receptor, C5aR2, we observed a partial, yet significant agonistic effect from SB290157. Albeit being less potent than the existing C5aR2-selective agonist P32 (29), SB290157 displayed an improved efficacy at C5aR2. This result indicates that higher doses of SB290157 could activate C5aR2 to modulate immune responses. Indeed, multiple studies have utilised P32 as a pharmacological tool to decipher the roles of C5aR2 (29, 36–38), and we recently documented that P32 exerts profound modulatory effects on multiple signalling and functional responses of HMDMs (23). The partial C5aR2-agonistic activity of SB290157 at higher concentrations could therefore have significant functional implications *in vitro* and *in vivo*, and should not be overlooked.

In a previous study by Proctor et al. (16), SB290157 demonstrated anti-inflammatory activities in a rat model of intestinal ischemia/reperfusion injury, which was associated with ligand-induced global neutrophil tissue sequestration during ischemia, rather than pure C3aR antagonism. C5aR2 is expressed on polymorphonuclear leukocytes, including neutrophils, and has been to shown to suppress C5a-induced neutrophil chemotaxis, both *in vitro* and *in vivo* (28, 29). Given the results of our study, it is possible the non-specific neutrophil effect of SB290157 depicted by Proctor et al. (16) could have resulted from the ligand-induced activation of C5aR2.

It is not uncommon for SB290157 to be used at doses higher than needed to block C3aR in *in vivo* studies. For example, doses of 10-30 mg/kg SB290157 are commonly used *in vivo* (12, 13, 39), and doses of 10 μM are used *in vitro* (40, 41). Based on prior pharmacokinetic reports as summarized in **Table 1**, intravenous (*i.v.*) administration of any dose greater than 1 mg/kg could result in a sufficient circulatory concentration of SB290157 to target both C3aR and C5aR2 based on its calculated IC_50_ and EC_50_. Likewise, intraperitoneal (*i.p.*) administration of dose greater than 10 mg/kg could also lead to a circulatory concentration that ultimately targets both receptors. Hence, administering a dose greater than 10 mg/kg (*i.p.*) or 0.3 mg/kg (*i.v.*), would likely result in both agonistic activity at C5aR2 and antagonist/agonist activity at C3aR. High *in vivo* doses of SB290157 could therefore lead to numerous on- and off-target adverse effects such as neutropenia (16), transient hypertension (16), gain of body weight (12), hematopoietic effects (42) and tachycardia (43), that would severely impact on data interpretation. Our data thus further highlights the importance of proper characterisation of drug pharmacokinetics to pharmacological profile when utilising compounds *in vivo* (17). Considering the broad expression of C5aR2 in immune cells and intricate interplay between C5aR2 and C3aR (4, 23, 44), the utilisation of high concentrations of SB290157 in primary immune cells, or *in vivo*, should be avoided whenever possible.

**Table 1:** Calculated molar concentration of SB290157 after various routes and doses of administration

| Species | Route | Dose (mg kg<sup>-1</sup>) | C<sub>max</sub> | Predicted C<sub>0</sub> | Molar concentration | Reference |
| --- | --- | --- | --- | --- | --- | --- |
| Rat | *i.v.* | 1 | ----- | >15 µg/ml | >36 µM | Proctor et al. (16) |
|  |  | 0.3 | ----- | >3 µg/ml | >7 µM |  |
|  |  | 0.1 | ----- | >1 µg/ml | >2 µM |  |
| Guinea pig | *i.p.* | 30 | >7 µg/ml | ----- | >16 µM | Ames et al. (10),<br>Woodruff et al. (17) |
| Mice | *i.p.* | 1; 3 times per week | >0.2 µg/ml | ---- | >400 nM | Lian et al. (14),<br>Woodruff et al. (17) |

Finally, despite the on- and off-target limitations of SB290157 (i.e. C3aR agonist and C5aR2 agonist activity), in human monocyte-derived macrophages that naturally express complement-peptide receptors, SB290157 effectively and potently inhibited C3a-induced phospho-ERK1/2 signalling, without any agonistic effect. This confirms prior reports that the pharmacological C3aR agonist activity of SB290157 may be dependent on cellular receptor density (15). As such, extra precaution needs to be taken when interpreting data generated with this ligand in overexpression cell systems, or native cells with naturally high levels of C3aR, such as mast cells (6), or under situations where there may be an upregulation of C3aR expression (45, 46). It should also be noted that we cannot exclude the possibility that some of SB290157-mediated dampening effects on C3aR-mediated ERK signalling in HMDMs indirectly resulted from C5aR2 agonism (23), especially when high concentrations of SB290157 were used

In conclusion, our results confirm that the widely used C3aR antagonist SB290157, exerts potent C3a agonist activty in transfected cells, and partial C5aR2 agonist activity at higher concentrations. As many studies continue to use this drug at concentration ranges that would exert activity at C5aR2, it is possible some of the reported phenotypes are due to activity at this receptor, in addition to any potential activity at C3aR. Thus, our data further caution against the use of this compound as the sole tool to delinate C3aR biology and function. Until further validated selective C3aR inhbitory tools become available (32), we strongly recommend the use of other genetic approaches such as gene-knockout mice, or gene knockdown approaches, to validate roles for C3aR. If SB290157 usage is absolutely required, an *in vivo* dose of no more than 1 mg/kg (*i.p.*) be considered to minimise off-target activity at C5aR2.

## Abbreviations used in this article

BRET: bioluminescence resonance energy transfer
BSA: bovine serum albumin
C3aR: C3a receptor
C5aR1: C5a receptor 1
CHO-C3aR: Chinese hamster ovary cells stably expressing C3aR
CHO-C5aR1: Chinese hamster ovary cells stably expressing C5aR1
DMEM: Dulbecco’s Modified Eagle’s Medium
ERK1/2: extracellular signal-regulated kinase 1/2
FBS: foetal bovine serum
HEK293: human embryonic kidney 293 cells
HMDM: human monocyte-derived macrophage
*i.p.*: intraperitoneal
*i.v.*: intravenous
rhC5a: recombinant human C5a
RT: room temperature
S.E.M.: standard error of the mean
SFM: serum-free medium

## 5 Conflict of Interests

All authors declare no conflict of interest pertaining to this manuscript.

## 6 Author Contributions

T. M. Woodruff, J. D. Lee and X. X. Li conceived the project and designed the research; X. X. Li performed the research, analysed data and wrote the first paper draft. V. Kumar provided additional pharmacokinetic data analyses and insight. All authors edited the manuscript, and approved the final version.

## 7 Funding

This work was supported by National Health and Medical Research Council of Australia (NHMRC) [Grant APP1118881 to TMW].

## 8 Acknowledgements

We would like to acknowledge Australian Red Cross Lifeblood and human donors for providing the cells used in these studies.

## Notes

### Competing Interest Statement

The authors have declared no competing interest.

